# Isoform-level profiling of m6A epitranscriptomic signatures in human brain

**DOI:** 10.1101/2024.01.31.578088

**Authors:** Josie Gleeson, Sachithrani U. Madugalle, Catriona McLean, Timothy W. Bredy, Ricardo De Paoli-Iseppi, Michael B. Clark

## Abstract

The RNA modification N6-methyladenosine (m6A) is highly abundant in the human brain and implicated in neuropsychiatric and neurodegenerative disorders. However, most techniques for studying m6A cannot resolve modifications within RNA isoforms and we lack an isoform-level map of m6A sites in the brain. Profiling m6A within isoforms is therefore a critical step towards understanding the complex mechanisms that underpin brain function and disease. Oxford Nanopore direct RNA sequencing (DRS) can quantify isoform expression, modifications and polyA tail lengths, enabling simultaneous investigation of the transcriptome and epitranscriptome. We applied DRS to three post-mortem human brain regions: prefrontal cortex, caudate nucleus and cerebellum. We identified 57,000 m6A sites within 15,000 isoforms and estimated that >27% of mRNA molecules contained an m6A modification. Our results revealed both isoform- and brain-region-specific patterning of m6A modifications and polyA tail lengths. The prefrontal cortex exhibited a distinctive profile of specifically modified isoforms enriched in excitatory neuron cell types and also had the highest proportion of previously unannotated m6A sites. A population of isoforms were hypermodified with m6A and were associated with excitatory neuron cell types in all three brain regions. We also discovered >15k differentially expressed isoforms, >2k differentially modified m6A sites and 566 isoforms with differential polyA lengths between brain regions. Our study demonstrates the utility of DRS for investigating multiple features of RNA isoforms in the brain and provides new insights into brain region specificity and functioning with implications for neurological development and disease.

## INTRODUCTION

Complex mechanisms of gene regulation are critical for the unique functioning and development of the human brain. A single gene can produce multiple RNA isoforms through alternative splicing and polyadenylation processes, greatly expanding the transcriptional diversity of both protein-coding and non-coding RNAs (1). Different gene isoforms commonly have distinct post-transcriptional fates and can encode RNAs and protein products with varying or opposing functions (2,3). The brain has the highest levels of splicing activity in human tissues and various neuronal pathways are regulated by differential expression of isoforms, such as cell fate determination, axon guidance, and synaptogenesis (4).

Post-transcriptional chemical modifications can also regulate the function of protein-coding and non-coding RNAs. The most abundant internal mRNA modification in eukaryotes is N6-methyladenosine (m6A), which regulates many aspects of the brain transcriptome (5–8). The brain has the highest levels of m6A in human tissues, which increases from developmental stages into adulthood (5,9). The dysregulation of RNA modification processes has been implicated in many neurodegenerative and neuropsychiatric disorders (10), and m6A is critical for brain development, learning and memory (11,12).

Due to the established importance of differential isoform expression in the human brain, it is essential to characterise m6A modification sites at the isoform level. Popular methods to study m6A involve immunoprecipitation of modified RNA fragments followed by short-read sequencing (SRS) (5,6). However, these methods only provide information on m6A modifications at the gene level. The exact nucleotide position and stoichiometry of m6A sites cannot be determined using these methods, and it is therefore often impossible to identify which original RNA isoform contained the modification (13,14). Chemical-based and enzyme-based detection methods that induce mutations at antibody binding sites enable the detection of m6A at single nucleotides (15,16). However, these techniques have had limited uptake as they do not provide isoform resolution and require complicated and expensive protocols. Therefore, there is a lack of knowledge about how m6A modifications are regulated at the isoform level, and it remains unknown whether isoforms are differentially modified within genes or between tissues.

Long-read direct RNA sequencing (DRS) from Oxford Nanopore Technologies (ONT) addresses many of these limitations by providing single-nucleotide isoform-level resolution of m6A modifications. DRS enables RNA sequencing without fragmentation or conversion to cDNA, preserving RNA modifications and polyA tail lengths. Additionally, the quantification of m6A modification rates with DRS is highly similar to that of enzymatic approaches (16–18). However, no studies have applied DRS to the human brain to investigate the critical role of m6A modifications in this complex organ.

We aimed to characterise isoform-level m6A modification sites across the human brain transcriptome and integrate this with both isoform expression and polyA tail lengths in different brain regions. To our knowledge, we have performed the first application of DRS to the human brain, profiling tissues from three functionally distinct regions: prefrontal cortex, caudate nucleus and cerebellum. We provide an isoform-level transcriptome-wide map of m6A modification sites and identify widespread changes in isoform expression, m6A profiles and polyA lengths both between gene isoforms and between the different brain regions. Our study reveals brain-region-specific regulation of m6A modifications within isoforms and shows that many specifically modified isoforms are associated with distinct cell types in different brain regions. We show that modification rates of m6A sites in different isoforms from a single gene are influenced mainly by isoform structure and proximity to downstream exon boundaries. Based on our findings, we recommend that m6A modifications be interpreted in isoform- and tissue-specific contexts.

## RESULTS

### Long-read direct RNA sequencing of human brain samples

We applied direct RNA sequencing (DRS) to post-mortem human brain samples from three brain regions: prefrontal cortex (PFC), caudate nucleus (CN) and cerebellum (CB) (Figure 1). DRS generated >52 million high-quality reads (qscore >7) from 10 samples with a median read length of 720 nt (Table 1). We included synthetic SIRV spike-in RNAs as a control and sequenced 360,695 SIRV reads. We identified the expression of >22k genes and >62k isoforms across the brain regions, and the reads covered a median of 59.50% of their mapped transcript isoform with a median accuracy of 90.82% (Supplementary Figure 1).

**Table 1.** Direct RNA sequencing metrics of 10 post-mortem human brain samples from 4 individual donors.

|  | PFC 1 | PFC 2 | PFC 3 | PFC 4 | CN 2 | CN 3 | CN 4 | CB 1 | CB 2 | CB 4 | Total |
| --- | --- | --- | --- | --- | --- | --- | --- | --- | --- | --- | --- |
| **Total pass reads (per million)** | 5.67 | 5.05 | 4.08 | 6.04 | 6.73 | 5.44 | 5.56 | 5.78 | 4.88 | 3.18 | 52.4 |
| **Median read alignment length (nt)** | 656 | 792 | 803 | 656 | 643 | 687 | 580 | 753 | 919 | 756 | 720 |
| **Median coverage fraction of reads** | 0.43 | 0.60 | 0.62 | 0.46 | 0.56 | 0.59 | 0.48 | 0.61 | 0.69 | 0.61 | 0.59 |
| **Median accuracy of reads (%)** | 90.30 | 91.10 | 90.59 | 91.23 | 91.05 | 90.20 | 90.89 | 90.83 | 90.81 | 90.31 | 90.82 |
| **Number of genes identified (count>5)** | 15,311 | 14,815 | 14,052 | 15,051 | 14,989 | 14,921 | 15,025 | 15,077 | 14,786 | 13,389 | 22,608 |
| **Number of isoforms identified (count>5)** | 33,501 | 31,420 | 28,534 | 32,890 | 32,055 | 31,271 | 32,032 | 33,153 | 32,078 | 27,104 | 62,577 |

**Figure 1.**
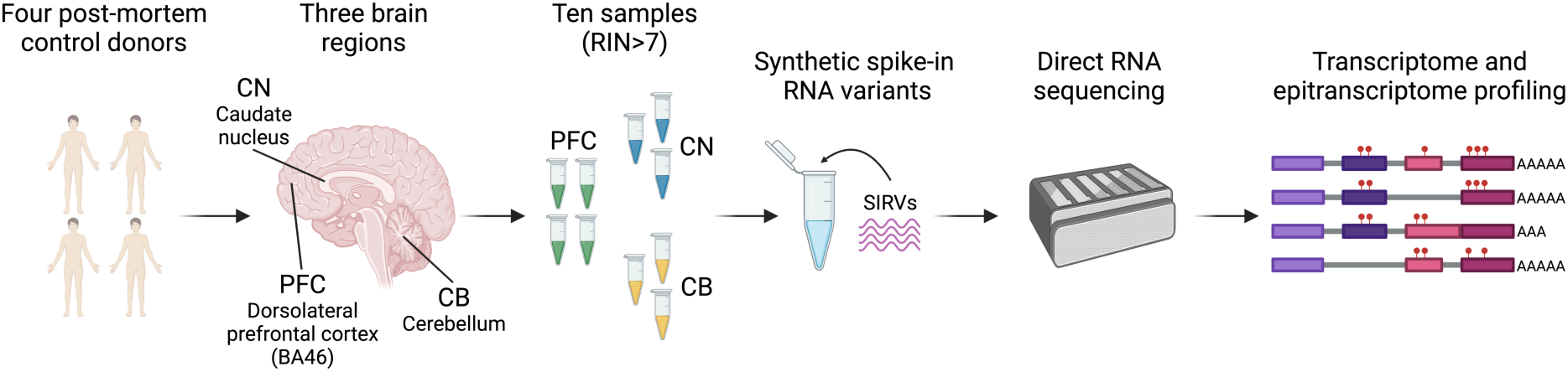
Experimental overview of direct RNA sequencing of post-mortem human brain samples. RNA was isolated from brain tissue of donors without neurological disorders from: prefrontal cortex (PFC), caudate nucleus (CN), and cerebellum (CB) (Supplementary Table 1). Samples with an RNA integrity score (RIN) >7 were sequenced using ONT’s PromethION device. Spike-in RNA variants (SIRVs) were added as controls.

### Identification of brain-region-specific transcriptional patterns and isoform switches

We explored expression differences between the brain regions and found that samples clustered by brain region for both gene and isoform expression, with CB having the most distinct expression profile compared to both PFC and CN (Figure 2A,B). We found ten thousand (n=9,908) differentially expressed genes (DEGs) between the brain regions (Figure 2C, Table 2, Supplementary Table 2), many of which confirmed previous observations of genes known to be upregulated in particular brain regions, such as increased expression of *DRD2* in CN (19).

**Figure 2.**
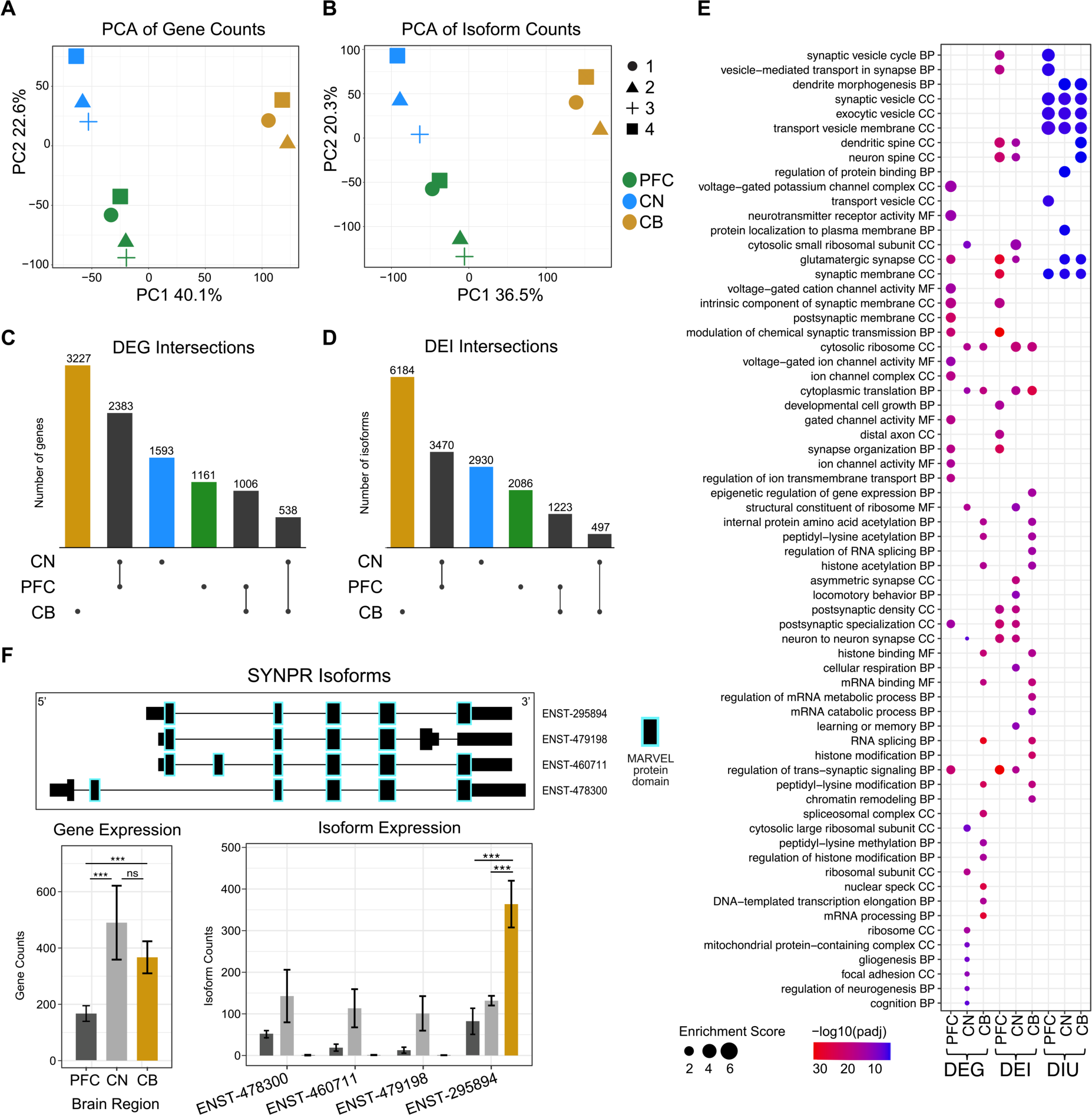
Widespread gene and isoform expression differences between brain regions. Principal component analysis (PCA) of **(A)** gene counts and **(B)** isoform counts from human brain samples. Colours indicate brain regions (PFC=green, CN=blue, CB=yellow) and shapes indicate individual donors. **(C)** UpSet plot of differentially expressed genes (DEGs) between brain regions. **D.** UpSet plot of differentially expressed isoforms (DEIs) between brain regions. **(E)** Gene ontology (GO) analysis of DEGs, DEIs and genes with DIU in each brain region. **(F)** *SYNPR* gene and isoform expression plot. Isoform IDs have ‘-’ representing five 0s. Top panel shows the isoform structure with protein domains highlighted. Gene and isoform expression bar plots are shown below. In CB, only one isoform (ENST00000295894) was expressed whereas in PFC and CN, the gene expression is composed of multiple isoforms. Significance of adjusted p-values is indicated by ‘*’ for <0.05 and ‘***’ for <0.001.

**Table 2.** Genes and isoforms with significantly upregulated expression or usage in each brain region.

|  | DEG | DEI | DEI,<br>no DEG | Gene<br>with DIU | DIU | Gene with DIU,<br>no DEG | DIU,<br>no DEI |
| --- | --- | --- | --- | --- | --- | --- | --- |
| **PFC** | 4,550 | 6,779 | 1,328 (19.59%) | 199 | 251 | 121 (60.80%) | 71 (28.29%) |
| **CN** | 4,514 | 6,897 | 1,647 (23.88%) | 300 | 488 | 214 (71.33%) | 132 (27.05%) |
| **CB** | 4,771 | 7,904 | 2,029 (25.67%) | 310 | 346 | 199 (64.19%) | 83 (23.99%) |

We also identified 16,390 differentially expressed isoforms (DEIs) between brain regions (Table 2, Supplementary Table 2), and most DEIs displayed brain-region-specific upregulation. Isoforms upregulated specifically in CB were the largest category of DEIs, followed by those upregulated in both PFC and CN (Figure 2C,D). For example, the gene *SYNPR* encodes a synaptic vesicle component, synaptoporin, and was expressed in all three brain regions. However, only one isoform contributed to the expression profile in CB, whereas four isoforms were expressed in PFC and CN (Figure 2F).

Changes in the proportion each isoform contributes to gene expression between tissues are also biologically relevant and were examined with a differential isoform usage (DIU) analysis. The results of differential isoform expression (DIE) may largely reflect those of gene expression, which can mask complexity at the level of isoform usage. We found 764 isoforms encoded by 317 genes with differential usage between brain regions (Table 2, Supplementary Table 2). Of the features with DIU in each brain region, 26% of the isoforms and 65% of the genes did not have upregulated DE, highlighting the additional insight provided with DIU. GO analysis of genes with DIU identified a dramatically different profile to DEGs or DEIs (Figure 2E, Supplementary Table 3). The genes with DIU displayed a consistent signal for synapses and synaptic vesicles, suggesting a specific gene regulatory program for these genes involving isoform switching.

We identified several genes implicated in neurodevelopmental and neuropsychiatric conditions that exhibited DIU, including *PRMT7*, *RBFOX1* and *GRIA1* (20,21). The overall expression of *PRMT7* mRNA was highest in CN (Figure 3A). However, protein expression data from the Human Protein Atlas identified the opposite result (22). *PRMT7* DIU analysis revealed that PFC and CB both expressed a greater proportion of the canonical protein-coding isoform compared with CN and that most of the gene expression in CN was due to the expression of a shorter non-coding nonsense-mediated decay (NMD) isoform. This result highlights how expression at the gene level can mask underlying complexity at the isoform level.

**Figure 3.**
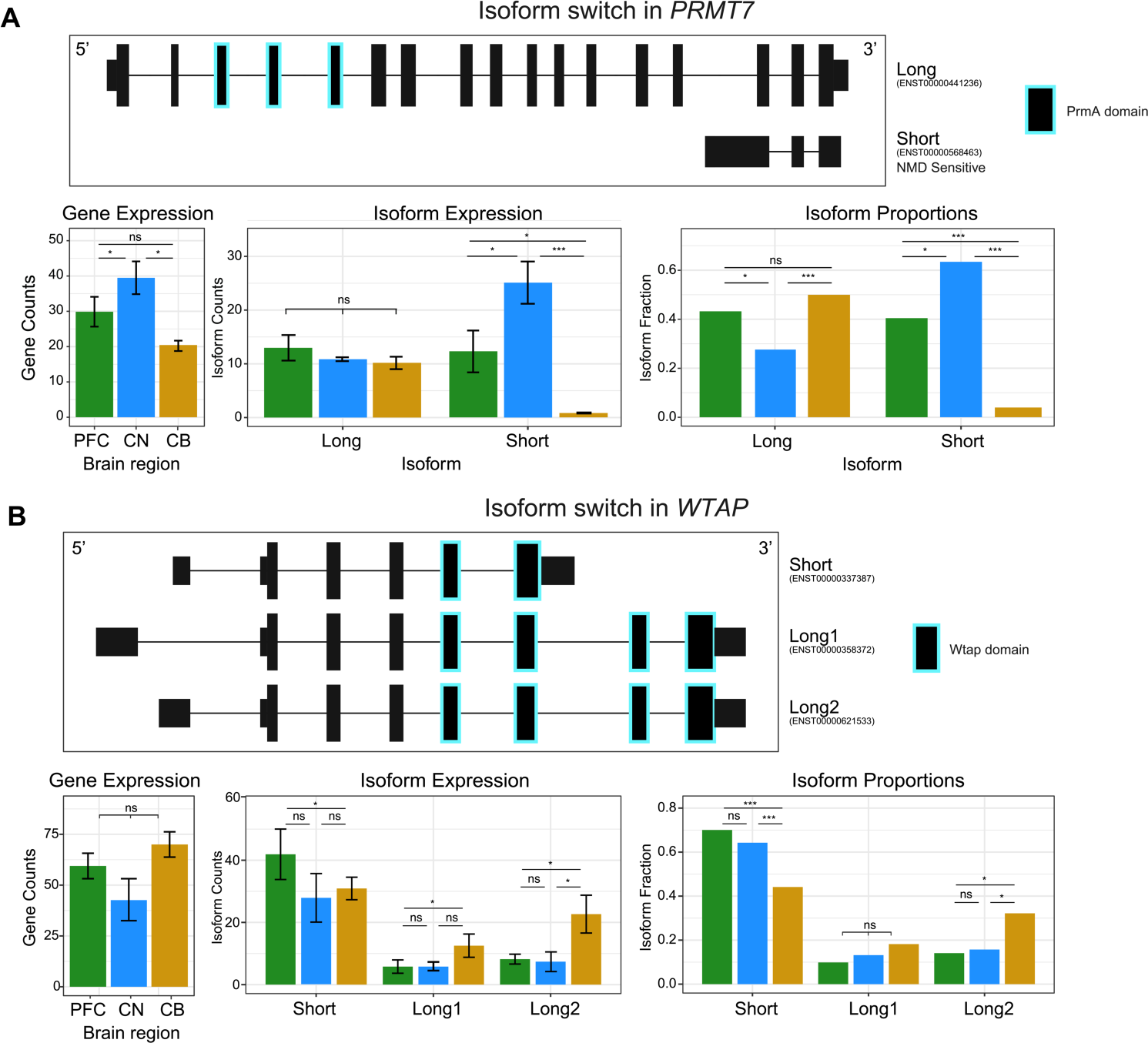
Clinically relevant genes exhibit different isoform usage between brain regions. Switch plots for isoforms with differential usage from **(A)** *PRMT7* and **(B)** *WTAP*. Top of each panel shows the isoform structures with protein domains highlighted. Gene expression, isoform expression and isoform proportions are shown as bar plots below for PFC (green), CN (blue) and CB (yellow). Significance of adjusted p-values is indicated by ‘*’ for <0.05 and ‘***’ for <0.001.

The *WTAP* gene, a subunit of the m6A writer complex, had an isoform switch in CB compared to PFC and CN (Figure 3B). WTAP interacts with METTL3, METTL14 and VIRMA to control m6A modification levels on RNA. The majority of expression in CB was from two longer isoforms that contained the complete WTAP protein domain. In contrast, PFC and CN primarily expressed a short WTAP isoform missing two exons required for the WTAP protein to bind VIRMA (23,24). Despite no significant change at the gene expression level, this isoform switch may result in decreased activity of the m6A writer complex in PFC and CN.

### Isoform-level map of m6A modification sites in the human brain

DRS enables the identification of m6A modification sites at the isoform level with single nucleotide resolution, allowing us to determine the exact transcriptomic position of a modification and the modification rate (proportion of modified reads) at these sites using m6anet (18). We tested 1.14 million DRACH sites for m6A modification, identifying 73,843 sites with an m6A modification probability of >0.9. We further filtered these for sites reported as modified in >1 sample, resulting in 57,144 high-confidence m6A sites (Methods) (Supplementary Table 4). All downstream analysis was performed on these high-confidence sites. We also tested for m6A modifications within the unmodified SIRV control reads and did not identify any m6A sites. The m6A sites followed a typical distribution with enrichment around stop codons and in 3’ untranslated regions (UTRs) (5,6) (Figure 4A).

**Figure 4.**
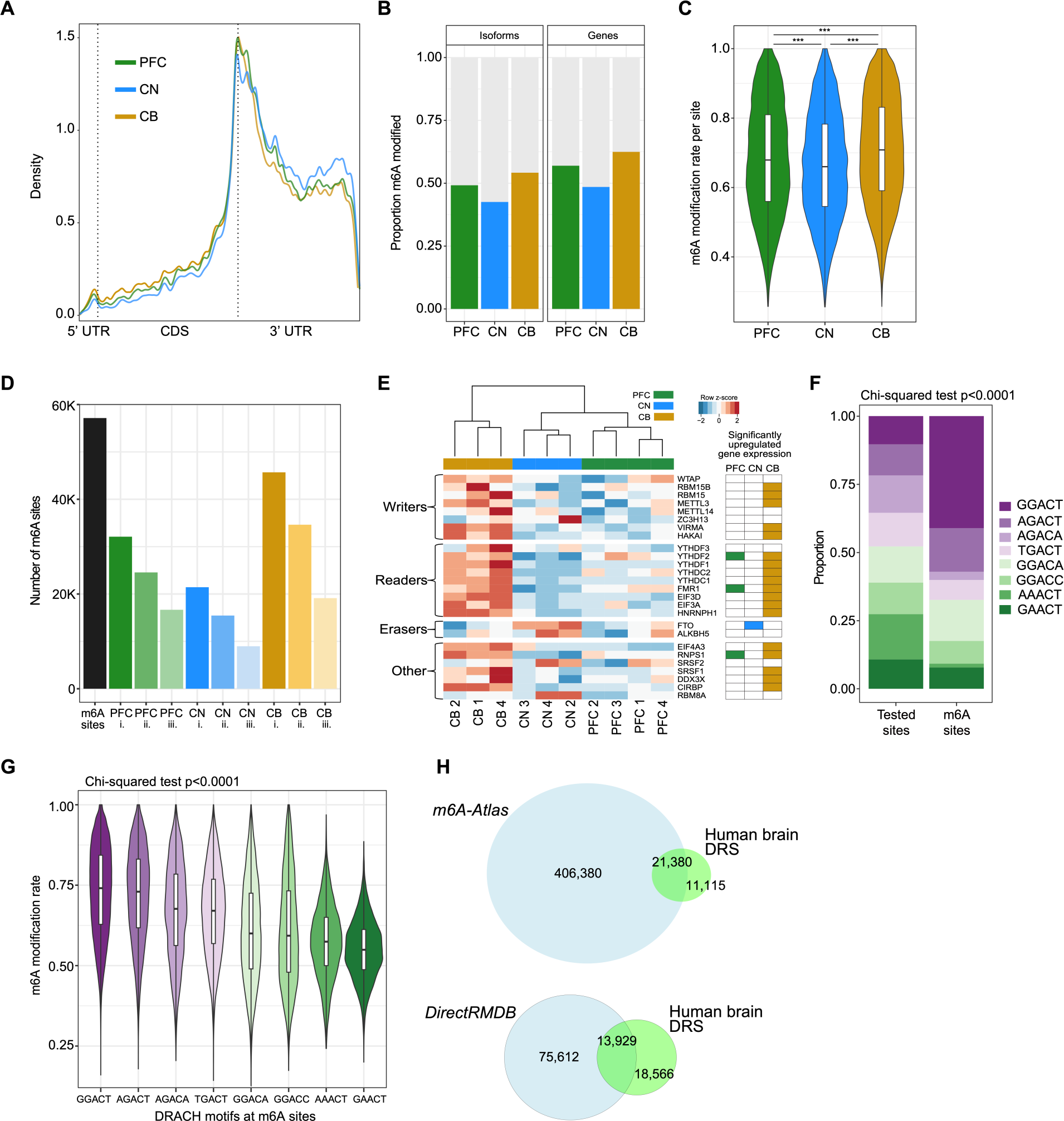
Isoform-level m6A modification sites in three human brain regions. **(A)** Metagene plot showing the distribution of m6A modification sites along mRNA isoforms for PFC, CN and CB. **(B)** Proportion of modified genes and isoforms in each brain region. **(C)** Modification rate per m6A site in each brain region. **(D)** Number of m6A sites detected in >1 brain sample (n=57,144) (black). M6A sites per brain region detected in **i.** ≥1 sample, **ii.** ≥2 samples, and **iii.** ≥3 samples. **(E)** Heatmap of gene expression of m6A-related genes across three brain regions. **(F)** Proportions of each DRACH motif in all tested sites and m6A sites. **(G)** Modification rates per DRACH motif for all m6A sites. **(H)** Venn diagrams showing intersections of m6A sites identified in our human brain DRS data compared with those identified in either *m6A-Atlas* or *DirectRMDB* (27,28).

There were 15,368 isoforms modified with m6A across the brain regions and a mean of three m6A sites per isoform, consistent with previous studies (5,6). More than half of the total detected genes (65%) and isoforms (55%) were modified with m6A (Figure 4B), and m6A sites had a median modification rate of 0.66 (Figure 4C). We estimated that at least 27% of the >30 million reads tested for modifications contained at least one m6A site. These results demonstrate that many mRNA molecules are m6A modified in human brain tissue and that these sites are commonly found with high stoichiometry.

We assessed the proportion of m6A sites identified in ≥3 samples per brain region and found that approximately 45% met this threshold (Figure 4D). We compared the coverage (number of reads) and modification rate of m6A sites detected in ≤2 vs ≥3 samples per brain region. We found that the latter had significantly higher coverage (mean read coverage increase >15 reads, Mann-Whitney-Wilcoxon (MWU) p<0.0001) and modification rates (mean rate increase >20%, MWU p<0.0001) (Supplementary Figure 2A,B). Therefore, read coverage and modification rates both affect the reproducibility of m6A sites between samples and higher reproducibility would be obtained with higher read depths per sample. We also found that the modification rates at m6A sites were highly correlated between samples from the same brain region (Supplementary Figure 2C).

Cerebellum had the highest percentage of modified genes (63%) and isoforms (54%), as well as the highest median modification rate (0.71), consistent with previous studies of CB in mice (25,26). Expression of multiple genes encoding m6A writers and readers was significantly higher in CB, which may account for the increased levels of m6A modification observed in this brain region, while the m6A eraser *FTO* was significantly upregulated in CN (Figure 4E) (Supplementary Table 5). Notably, samples clustered by brain region even when subsetting expression to only m6A-related genes, suggesting that brain region differences in m6A profiles may partly be due to region-specific regulation of the m6A machinery.

It has been previously observed that high m6A modification levels are associated with long 3’ UTRs of isoforms (29,30). We investigated whether isoforms in our data had changes in 3’ UTR lengths in the different brain regions that were contributing to some of the differences in m6A modification levels. We compared both the total isoform lengths or 3’ UTR lengths between each brain region and found no differences in these features in all detected isoforms (counts >5). However, when we compared the isoforms that were upregulated (DIE) or m6A modified in each brain region, there were significant differences in both the total length and 3’ UTR lengths between brain regions (Supplementary Figure 3A-C). CB had longer isoform and 3’ UTR lengths than CN in both cases, consistent with the increased modification levels observed in CB. Therefore, the upregulation of longer isoforms in CB may partially drive the increased levels of m6A in CB, and the differences in modification levels observed between brain regions are, in part, a consequence of tissue-specific isoform expression patterns.

Examination of DRACH motifs revealed ‘GGACT’ was the most commonly modified motif, significantly enriched compared to its abundance within RNA, while ‘GGACT’ and ‘AGACT’ had the highest modification rates (16,31) (Figure 4F,G). No correlation was observed between the modification rate at m6A sites and the motif frequency in m6A sites. All brain regions harboured similar proportions of each DRACH motif and the m6A sites in CN had consistently lower modification rates compared with the other brain regions overall as well as within each DRACH motif (Figure 4C, Supplementary Figure 3D). Therefore, we expect that the lower m6A levels observed in CN are likely due to differences in the expression of m6A-related genes and the expression of particular isoforms rather than a bias towards specific motifs.

### Discovery of previously unannotated m6A modification sites

We compared our gene-level m6A modification sites (n=29,596) with those previously annotated in two m6A databases, *m6A-Atlas* or *DirectRMDB* (27,28), and found that 71.36% of the sites in our data had been previously annotated in human tissues or cell lines (Figure 4H). The unannotated sites in our data had only marginally lower modification probabilities (-0.0159, p<0.0001) and rates (-0.0021, p<0.0001) compared with the annotated sites. We found that long non-coding RNA (lncRNA) biotypes were enriched in unannotated sites in our data as well as *DirectRMBD* sites compared with those in *m6A-Atlas* (p<0.0001), suggesting that DRS may be a more suitable technique for identifying m6A within lncRNAs than previous methods (Supplementary Figure 4).

Although CB had the highest total number of m6A sites, PFC had the highest percentage of unannotated m6A sites (26.05%), as well as the highest percentage of genes found with only unannotated modification sites (17.41%). Hence, while the CB displayed a higher frequency of modification, the PFC exhibited a less annotated and more distinctive m6A modification profile. Genes not previously identified as m6A modified were associated with brain-specific gene ontology (GO) terms such as ‘regulation of synaptic plasticity’, ‘synaptic signalling’, and ‘behaviour’ (Supplementary Table 6). In contrast, genes with only known sites were associated with more general terms such as ‘RNA splicing’, ‘protein catabolic process’ and ‘cellular protein localisation’, which highlights the additional information on m6A modifications that DRS can provide, as well as the existence of previously unidentified m6A modifications on key brain genes (Supplementary Table 6).

### Common and brain-region-specific m6A modification of RNA isoforms

Of the >50k m6A modification sites identified in total, there were 5,257 and 22,930 identified in all ten samples or all three brain regions, respectively. Although a majority (67%) of modified isoforms (n=10,253) were modified in multiple brain regions, 33% were only modified in a single brain region (n=5,115 isoforms) (Figure 5A). We integrated the results from our DE analysis and found that most of the genes and isoforms modified in only a single brain region were not uniquely expressed or specifically upregulated in those brain regions (Table 3, Supplementary Table 7). Therefore, region-specific m6A modification was not simply due to region-specific expression (25).

**Figure 5.**
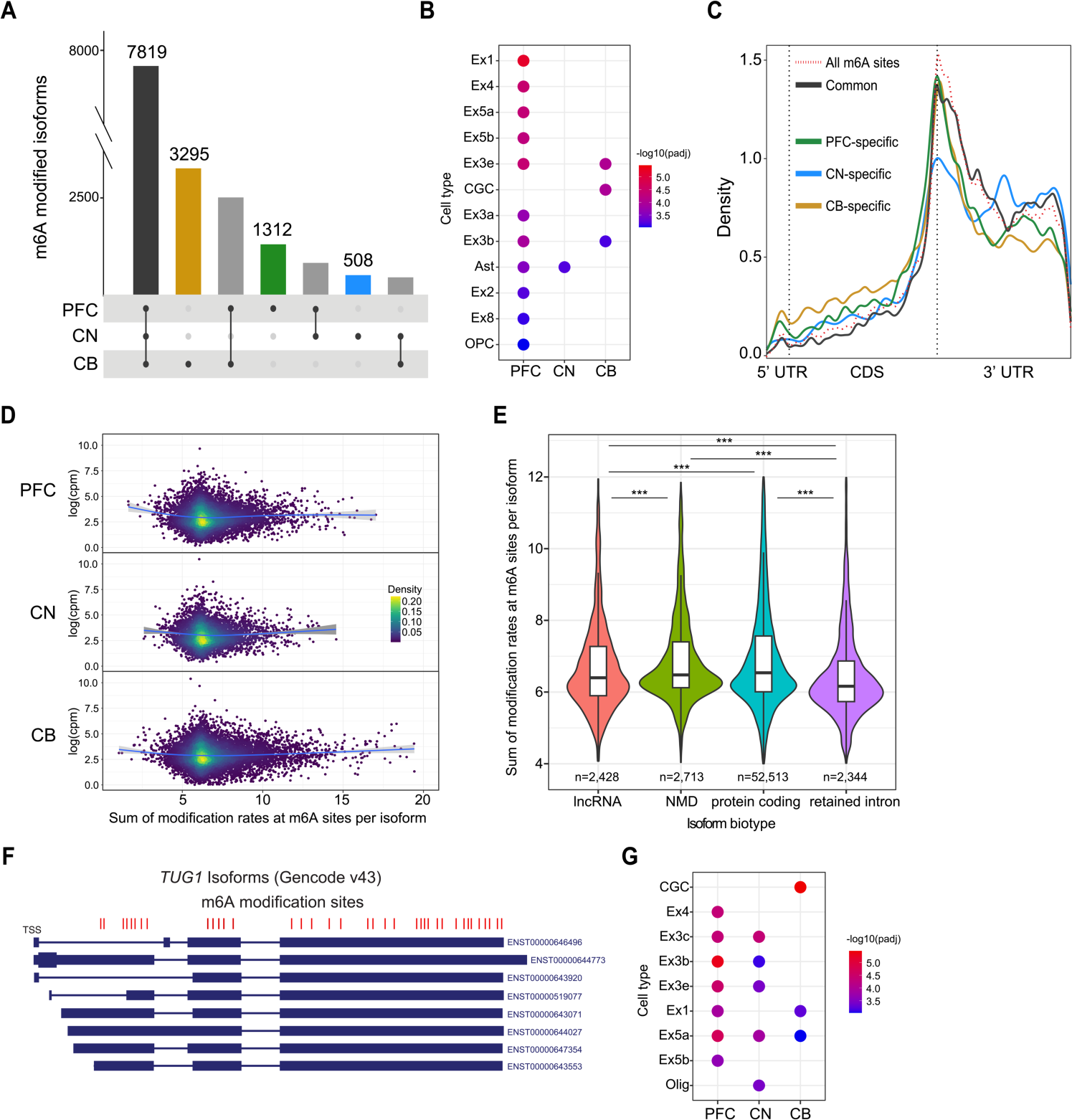
Brain-region-specific m6A modification patterns. **(A)** Isoforms with common and specific m6A modification in each brain region. **(B)** Cell-type-specific analysis of the genes encoding isoforms with specific m6A modification (and no expression upregulation) in each brain region. Ex=excitatory neuron, CGC=cerebellar granule cell, OPC=oligodendrocyte precursor cell, Ast=astrocyte. **(C)** Metagene plot showing the distribution of m6A modification sites specific to each brain region and common to all brain regions. The background of all m6A modification sites is shown with the dashed red line. **(D)** Scatter plots coloured by density for expression in log(counts per million) per isoform compared to summed modification rates at m6A sites per isoform for each brain region. **(E)** Summed modification rates at m6A sites per isoform plotted per transcript isoform biotype. Only the significant MWU comparisons are labelled. Protein-coding vs retained intron p<0.0001; Protein-coding vs lncRNA p<0.0001; Nonsense-mediated decay vs retained intron p<0.0001; Nonsense-mediated decay vs lncRNA p<0.0001. **(F)** Isoforms of the most highly modified gene, *TUG1*. M6A modification positions are shown in red. **(G)** Cell-type-specific analysis of the genes encoding the top 500 most highly modified isoforms in each brain region. Ex=excitatory neuron, CGC=cerebellar granule cell, Olig=oligodendrocyte.

**Table 3.** Specific m6A modification and expression of genes and isoforms in the different brain regions.

|  | Region specific m6A modification<br>(gene / isoform) | Region specific DE upregulation<br>(gene / isoform) | Region specific m6A, no DE upregulation<br>(gene / isoform) | Region specific m6A, no DE upregulation %<br>(gene / isoform) |
| --- | --- | --- | --- | --- |
| **PFC** | 553 / 1312 | 1161 / 2086 | 449 / 1065 | 81% / 81% |
| **CN** | 226 / 508 | 1593 / 2930 | 145 / 322 | 64% / 63% |
| **CB** | 1413 / 3295 | 3227 / 6184 | 790 / 1953 | 56% / 59% |

The CB had the largest number of specifically modified features and the largest proportion likely due to increased expression. The PFC exhibited the highest proportion (81%) of modified genes and isoforms not due to expression differences, underscoring the distinctive regional regulation of m6A modification in the PFC (Table 3). We performed a GO analysis on isoforms with region-specific m6A modification (without region-specific expression) and isoforms commonly modified in all three brain regions. The commonly modified isoforms were primarily associated with protein catabolic terms and, secondarily, neuronal and synaptic terms. CB-specific and CN-specific isoforms had no associations, whereas PFC-specific m6A isoforms were associated with multiple synaptic and neuronal cellular components (Supplementary Table 8).

We also performed a cell-type-specific enrichment analysis to investigate whether the specifically modified isoforms were associated with different cell types in the different brain regions (32). PFC-specific m6A isoforms showed the highest degree of enrichment for multiple cell types, including for multiple subtypes of excitatory neurons (Figure 5B) (Supplementary Table 8). The CN-specific isoforms were only enriched for astrocytes, and the CB-specific isoforms were enriched for both cerebellar granule cells (CGCs) and two excitatory neuron subtypes. The integration of our DE analysis with the m6A modification data suggests that region-specific modification is spatially regulated by mechanisms other than isoform expression and that there are cell-type-specific roles of m6A in different brain regions. The cell types associated with specifically modified isoforms are consistent with differences in cell type composition across brain regions driving isoform-specific modification.

We also found that brain-region-specific modification sites had different distributions along an isoform than the common modification sites in all brain regions. The brain-region-specific modification sites in PFC and CB had increased densities in the 5’ UTR and CDS, and decreased densities in the 3’ UTR, compared to the common sites (Figure 5C, Kolmogorov–Smirnov p<0.0001). The observed divergence in distributions underscores the region-specific regulation of m6A modifications.

### Hypermodified and unmodified isoforms

We found that the total number of m6A sites per isoform was positively correlated with isoform length (rho=0.2163, p<0.0001), 3’ UTR length (rho=0.1289, p<0.0001) and negatively correlated with exon density (isoform length/number of exons) (rho=-0.2981, p<0.0001) (33,34). In agreement with recent studies, unmodified isoforms (n=2,339) isoforms were generally shorter in length with a higher exon density compared with modified isoforms (Supplementary Figure 5A,B) (MWU p<0.0001) (33,34). We normalised for isoform length and exon density to rank isoforms based on their overall modification levels (Methods). There was no correlation between the normalised number of m6A sites (or raw number of m6A sites) and isoform expression (Figure 5D). However, different transcript isoform biotypes had minor changes in modification levels. Protein-coding and nonsense-mediated decay (NMD) isoforms had higher m6A levels than retained intron (RI) and lncRNA isoforms (Figure 5E).

We found 911 hypermodified isoforms (616 genes) (Methods) and 413 isoforms were consistently hypermodified in multiple brain regions (45.33%) (Supplementary Table 9). The top hypermodified isoform in both PFC and CN was from the *PAQR8* gene (ENST00000360726), and in CB was from the *TUG1* lncRNA (ENST00000643071). Interestingly, *PAQR8* was also a top hypermethylated gene in the synaptic compartment of mouse forebrains (35). The *TUG1* lncRNA had 37 m6A sites, the highest total number observed on a gene in our data (Figure 5F) and most of these sites were found in all three brain regions. This lncRNA has been associated with glioma stem cell renewal and tumorigenesis, and it has been suggested that lncRNAs may regulate tumour growth through m6A modification (36,37). A study using SCARLET to profile m6A in lncRNAs tested ten sites in *TUG1* for the presence of m6A in HeLa, HEK293T and HEPG2 cell lines (38). However, only one site in *TUG1* was m6A modified, identified in all three tested cell lines. A more recent study investigated the m6A profile of *TUG1* in two glioma stem cell lines and identified nine m6A peaks across the gene (39). The variation in m6A modification patterns of *TUG1* across different studies and cell lines highlights a need for further research into the regulatory role of m6A modification of *TUG1* and other clinically relevant lncRNAs.

The hypermodified isoforms showed enrichment for excitatory neurons in all three brain regions and were highly associated with multiple synapse GO terms as well as ‘learning or memory’ and ‘cognition’ (Supplementary Figure 5C, Supplementary Table 10). The PFC hypermodified isoforms were exclusively enriched for excitatory neuron subtypes, whereas the CN and CB isoforms were also enriched for oligodendrocytes and CGCs, respectively (Figure 5G) (Supplementary Table 10). In contrast, unmodified isoforms were associated with cellular metabolism, respiration and ATP synthesis GO terms (Supplementary Figure 5D), Supplementary Table 10).

Hypermodified isoforms displayed increased modification density in the CDS compared to all modified isoforms and were not enriched for highly modifiable DRACH motifs (Supplementary Figure 5E,F). Previous work identified more CDS m6A sites amongst synaptic FMRP-target RNAs (25). The increased density of m6A in the CDS of hypermodified RNAs, coupled with their strong enrichment for synaptic processes and consistent association with excitatory neuron cell types, suggests the presence of a unique regulatory environment for synaptic RNA in excitatory neurons and that m6A modification has a key role in synaptic function.

### Differences in modification rates of sites between isoforms from the same gene

Recent studies have established that the exon junction complex and polyadenylation/transcription termination machinery binding creates an m6A modification exclusion zone of approximately 100 nt either side of splice junctions and transcription end sites (33,40). We assessed this in our m6A sites and found that distance to a downstream exon boundary was positively correlated with modification rate (rho=0.1731, p<0.0001) (Figure 6A). However, the distance to an upstream exon boundary was weakly correlated (rho=0.0256, p<0.0001).

**Figure 6.**
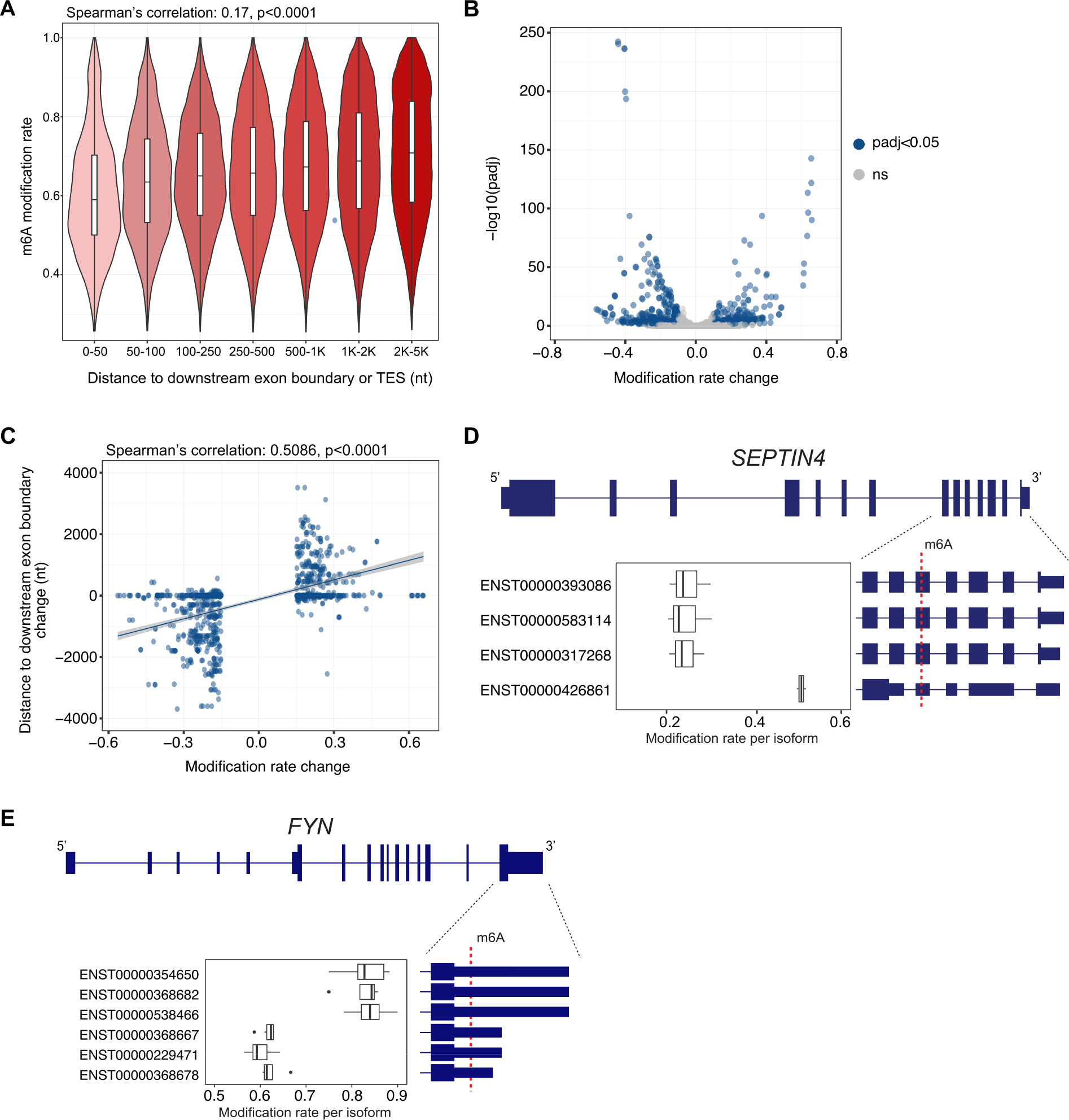
Differences in modification rates of sites between isoforms from the same gene. **(A)** The modification rate of m6A sites compared to the distance (in nucleotides, nt) to a downstream exon boundary or transcription end site (TES). **(B)** Volcano plot of modification rate differences within all tested genomic m6A sites (n=10,668) and corresponding p-values adjusted for false-discovery rate. Blue indicates significance (FDR<0.05) in a two-proportion z-test. **(C)** Modification rate differences of significant sites and corresponding changes in distance to a downstream exon boundary (or TES). **(D)** Genomic m6A site within *SEPTIN4* with significant differences in modification rate between isoforms (adjusted p-value<0.01). **(E)** Genomic m6A site within *FYN* with significant differences in modification rate between isoforms (adjusted p-value<0.05).

Building on this, we asked whether isoforms from the same gene could have different modification rates at the same genomic m6A site and, if so, whether these differences were associated with changes in isoform structure. We tested 10,668 genomic m6A sites encoded in 11,512 isoforms for differences in modification rates, and of these, 828 isoforms had significant differences (two-proportion z-test, FDR<0.05) at 320 differentially modified (DM) genomic sites (Figure 6B,C) (Supplementary Table 11). We found that an increase in modification rate in an isoform was strongly correlated with an increase in m6A site distance to a downstream exon boundary (or transcript end) (rho=0.5086, p<0.0001) and moderately correlated with an increase in distance from an upstream exon boundary (or transcript start) (rho=0.2650, p<0.0001). Most of the DM sites (n=264) had a change of >20 nt in the distance to an exon boundary between isoforms, while 30% of the total sites tested (n=3,208) had a change of >20 nt in the distance to an exon boundary between isoforms. Therefore, isoform structure is the main driver for isoforms with DM genomic m6A sites. However, most genomic sites shared between isoforms are in regions with consistent exonic structures.

Considering there were DM sites without changes in distances to an exon boundary, we investigated whether the location within the transcript region (5’ UTR, CDS, 3’ UTR) could also affect the modification rate. We included this in a linear regression along with the change in distance to an upstream or downstream exon boundary to predict modification rate differences. The model was highly significant (R^2^=0.4092, p<0.0001) and distances to exon boundaries were the most significant variables (upstream p<0.01, downstream p<0.0001). However, we found that genomic sites in 3’ UTRs had higher modification rates than the same sites within CDSs (p<0.01). For example, a DM site in *SEPTIN4* in CB had no differences in distance to exon boundaries between isoforms. However, the 3’ UTR site still had higher modification rates than the sites within the CDS (Figure 6D).

There were six isoforms encoding a DM site within *FYN* isoforms that showed the commonly observed pattern of increased distance to a downstream exon boundary and increased modification rate (Figure 6E). Three isoforms had long 3’ UTRs (1,407 nt), and three had short 3’ UTRs (457 nt). In both PFC and CN, the isoforms with long 3’ UTRs had an increase (mean=0.22) in m6A modification rates. Our single-nucleotide isoform-level m6A data allows an unbiased view of how mRNA structure affects modification by comparing the same position between different isoforms. Along with finding that regulation of m6A deposition can occur in an isoform-specific manner, our DM results demonstrate how proximity to splice junctions is not the only cause of differences in modification rates, which are also impacted by the distance to transcript ends and the CDS vs UTR status of a nucleotide.

### Differences in modification rates of isoform sites between brain regions

We hypothesised that the same site within an isoform may display brain-region-specific differences due to expression differences in m6A machinery or cell type composition between the brain regions. Changes within these sites would not be due to different 3’ UTR lengths or proximities to exon boundaries, as the isoform tested is identical between brain regions. We used xPore (41) to identify transcriptomic sites with DM rates between brain regions and found 2,218 significant DM sites within 1,658 isoforms (992 genes) (Supplementary Table 12). The majority of DM sites had an increase in modification rates in CB (n=1,666, 75.11%), consistent with the overall levels of m6A that we observed in this brain region (Figure 7A). Isoforms with DM sites were associated with microtubule polymerisation, protein transport and regulation of neuron projection GO terms (Figure 7B, Supplementary Table 12).

**Figure 7.**
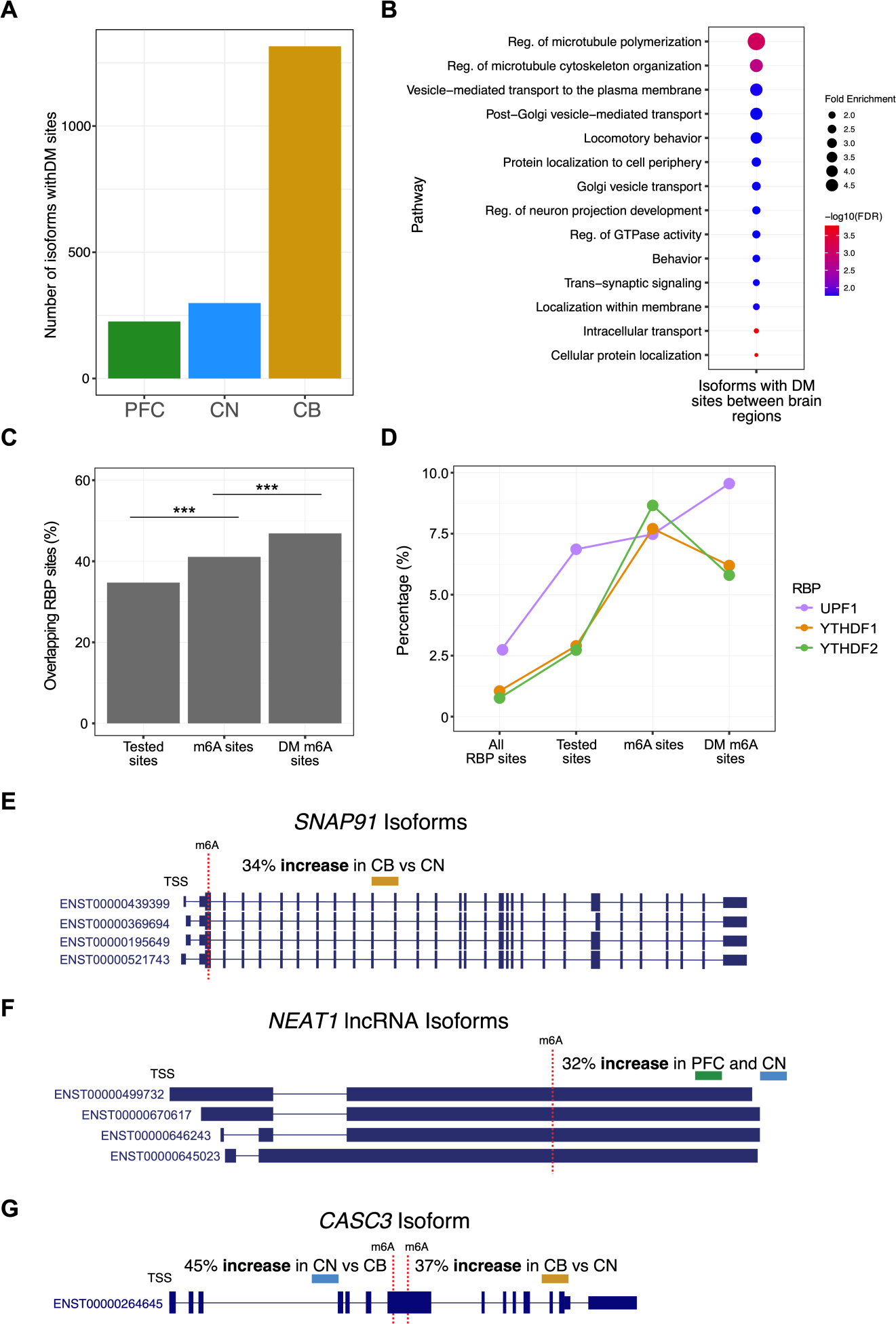
Differences in modification rates within transcriptomic m6A sites between brain regions. **(A)** The number of isoforms containing differentially modified (DM) m6A sites with increased modification rates in each brain region (n=1,658). **(B)** Gene ontology (GO) analysis of all gene isoforms with DM m6A sites. **(C)** The percentage of sites per category with genomic coordinates intersecting with RBP coordinates annotated in POSTAR3 and CLIPdb (43,44). Tested sites are DRACH sites tested for m6A modification, m6A sites are all m6A modified sites in m6anet and DM m6A sites are differentially modified sites identified by xPore. Two-proportion z-tests were performed for each comparison and were highly significant (‘***’ indicates p<0.001). **(D)** The percentage of sites in each category that overlap specified RBP binding sites, shown only for those with significant increases in either m6A sites or DM m6A sites. Two-proportion z-tests for: YTHDF1 tested sites vs m6A sites, YTHDF2 tested sites vs m6A sites, UPF1 m6A sites vs DM sites, were all significant with p<0.05. **(E)** An m6A site within four *SNAP91* isoforms with increased modification rates in CB compared to CN (p<0.05). **(F)** An m6A site within four *NEAT1* lncRNA isoforms with increased modification rates in PFC and CN compared to CB (p<0.05). **(G)** An m6A site within a *CASC3* isoform with opposing changes in modification rates in an internal exon. The proximal 5’ modification site had increased modification rates in CN compared to CB (p<0.001), whereas the distal 3’ site had increased modification rates (p<0.01).

In addition to gene expression changes of the m6A machinery in different brain regions, an additional explanation for DM of isoforms between brain regions could be due to changes in the activity of different RNA-binding proteins (RBPs) at these m6A sites. Previous studies have shown that highly m6A modified RNAs interact with more miRNAs and RBPs compared to unmodified RNAs (42). We intersected our data with RBP sites annotated in POSTAR3 and CLIPdb (Methods) (43,44). We found a striking enrichment for RBP sites in the DM m6A sites between brain regions compared with all m6A sites as well as all DRACH sites tested for m6A modification (Figure 7C) (43,45). The RBPs YTHDF1, YTHDF2 and UPF1 were significantly enriched in the DM sites compared with m6A sites. UPF1 had significantly increased expression in CB compared with the other brain regions in our gene expression data (p<0.0001) (and in GTEX) and directly interacts with the m6A reader protein YTHDF1 to promote rapid degradation of m6A modified RNAs (Figure 7D) (2,46). The enrichment of DM m6A sites for specific RBPs may highlight a set of isoforms that are spatially regulated between brain regions. The role of UPF1 in promoting RNA degradation may underscore that increased m6A levels in CB lead to increased RNA turnover rates, suggesting the importance of this mechanism in this particular brain region. Additionally, the enrichment of all m6A sites for the m6A readers YTHDF1 and YTHDF2 binding sites highlights the accurate identification of m6A sites in our dataset.

Four isoforms of *SNAP91* had sites with increased modification rates in CB compared to CN (Figure 7E). *SNAP91* is involved in synaptic function and is a risk gene for the development of schizophrenia (47,48). Little information exists about the role of m6A modification of *SNAP91*, though it is highly expressed in mouse Purkinje cells, suggesting this cell type likely drives this observation (49). In contrast, a site in *NEAT1* lncRNA isoforms displayed a ∼32% increase in m6A levels in PFC and CN compared to CB (Figure 7F). NEAT1 is essential for the formation of nuclear paraspeckles through extensive interactions with RBPs and is associated with neurodegenerative disorders (50,51).

Typically, isoforms with DM sites contained only one significant site. However, there were 153 isoforms with multiple DM sites and the majority of these exhibited a consistent change in direction with other DM sites in the same isoform across the brain regions. For example, two DM sites within the 3’ UTR of *NKAIN2* isoforms had a consistent increase in modification rates in CB compared to CN. Interestingly, there were 41 isoforms with inconsistent changes in the direction of modification rates. Two DM sites within a long internal exon (exon 7) of a *CASC3* isoform (ENST00000264645) had opposing changes in modification rates in CN and CB (Figure 7G). These results indicate that brain-region-specific regulation of m6A deposition occurs, which does not always follow the general trend observed in our data of increased modification levels in CB, demonstrating how additional factors likely control m6A modification at specific isoform sites.

### Regulation of isoform polyA lengths between brain regions

PolyA tails are critical in post-transcriptional regulation, including in stabilising mRNA and promoting translation (52). We used nanopolish (v0.13.3) to quantify the lengths of polyA tails in our human brain samples and tested for global changes in polyA tail length between brain regions as well as isoform-specific changes (53).

Samples from the same brain region mostly clustered together based on the median polyA length per isoform, demonstrating consistency between replicates (Supplementary Figure 6A). Samples from individual 3 were slightly more correlated with each other than samples in their respective brain regions. Globally, CN had shorter polyA tail lengths compared with both CB (-16 nt) and PFC (-13 nt) (MWU p<0.0001) (Figure 8A). The SIRV data showed no differences in polyA lengths between samples, brain regions or isoforms (Figure 8B). The polyA tails of SIRV isoforms had a median length of 35 nt, compared to the ground truth polyA length of 30 nt and polyA length estimates per SIRV isoform ranged from 27-46 nt.

**Figure 8.**
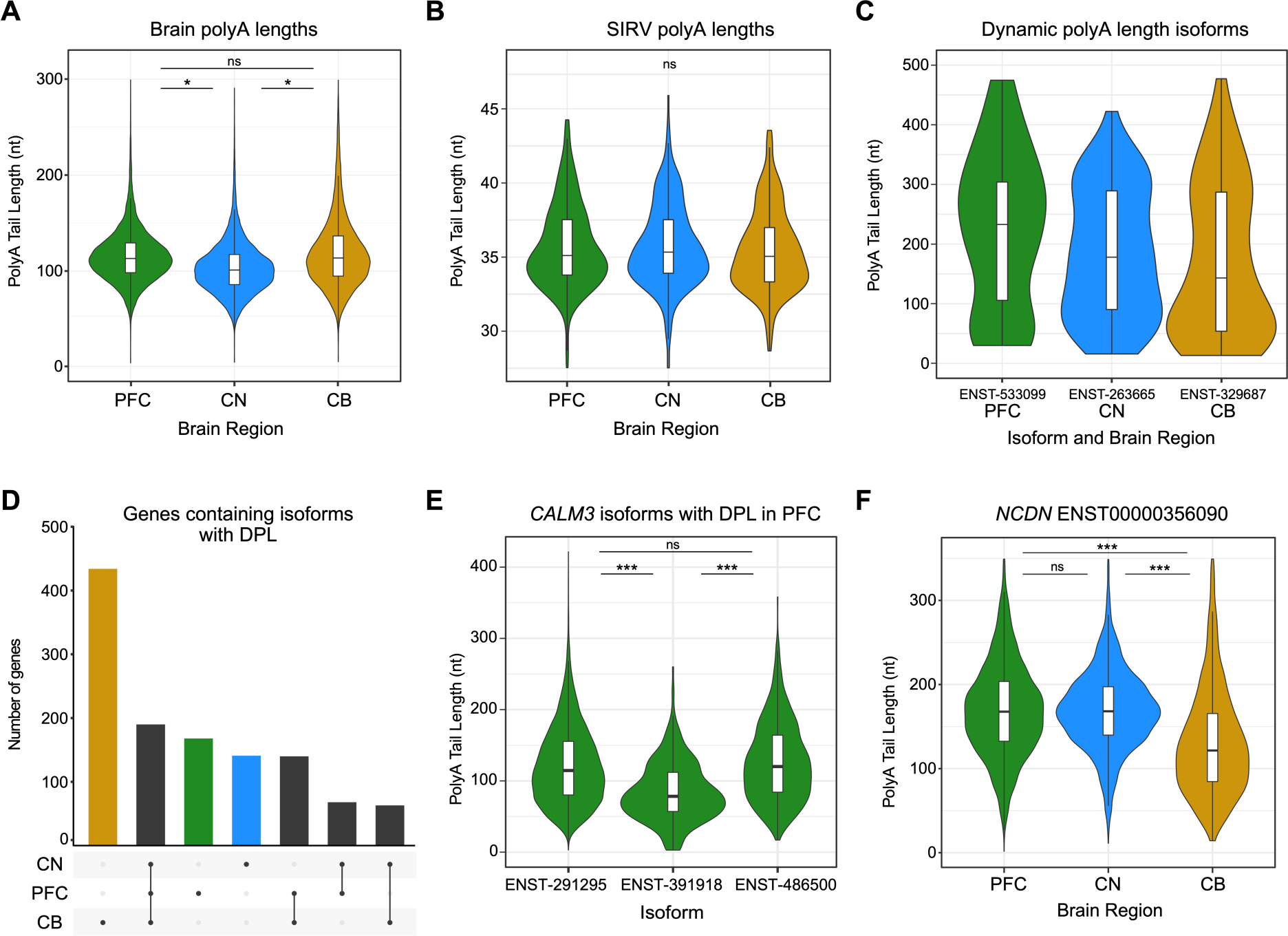
Quantification and changes in polyadenylation lengths between isoforms and brain regions. **(A)** Global polyA tail lengths per brain region plotted as a median value for every isoform in each sample. **(B)** Global polyA tail lengths of SIRV reads per brain region plotted as a median value for every isoform in each sample. **(C)** PolyA lengths of the most highly dynamic polyA isoform in each brain region. **(D)** UpSet plot of the gene isoforms found with differential polyadenylation lengths (DPL) within each brain region. **(E)** PolyA tail lengths of isoforms with DPL encoded by *CALM3* in PFC plotted for every read. **(F)** PolyA tail lengths of an *NCDN* isoform in each brain region were plotted for every read. Significance of p-values from Mann-Whitney-Wilcoxon tests are indicated by ‘*’ for <0.05 and ‘***’ for <0.001. Isoform ENST-IDs omit five 0s for clarity.

The median polyA length per isoform was positively correlated with annotated isoform length (rho=0.55, p<0.0001) and we did not observe significant differences in polyA lengths between different transcript isoform biotypes (Supplementary Figure 6B). We noted that some isoforms had considerable variation in their estimated polyA lengths, which we termed ‘dynamic’ polyA tails (Methods) (Supplementary Table 13). The most dynamic polyA length isoforms were from the genes *RNPC3*, *CNTN3*, and *GRIPAP1* in PFC, CN and CB, respectively, and these all had polyA tail length ranges of more than 400 nt (Figure 8C). The genes with dynamic polyA lengths were associated with RNA processing GO terms including RNA and ribonucleoprotein export, RNA localisation and RNA splicing (Supplementary Table 14).

We identified 3,545 isoforms encoded by 1,204 genes with differential polyA lengths (DPL) within genes (Supplementary Table 13). There were 190 genes that had isoforms with DPL in all three brain regions and a large portion of the genes were exclusively found with DPL in CB (Figure 8D).

We found that isoform length and 3’ UTR length were both positively correlated with polyA length (rho=0.5216, p<0.0001 and rho=0.4146, p<0.0001) and these factors were the main drivers of DPL within genes. For example, isoforms from *CALM3* had DPLs within the PFC (Figure 8E) and the isoform with the shorter polyA lengths (ENST00000391918) also had a short final exon. The genes from isoforms with DPL were associated with translation and splicing GO terms (Supplementary Figure 6C, Supplementary Table 14).

We next compared polyA lengths between the same isoform in different brain regions and identified 566 isoforms with differential polyA lengths (DPL) between brain regions (Supplementary Table 13). These mainly were between isoforms from CN compared to the other brain regions, reflecting the global trend of shorter polyA tails in CN. However, this was not always the case, as shown in an isoform from *NCDN* that had longer polyA tail lengths in both PFC and CN than CB (Figure 8F). The isoforms with DPL between brain regions were highly associated with synapse-related and neurotransmitter-related GO terms (Supplementary Figure 6D, Supplementary Table 14). These results suggest that polyA lengths of isoforms are likely under brain-region-specific regulation and that this process is particularly important in synaptic function.

### Integrative analysis of isoform expression, polyA tail lengths and m6A levels

We investigated whether isoform expression or m6A modification levels were related to polyA lengths and found a significant overlap (>1.7 times greater than expected by chance) between isoforms with increased polyA lengths and upregulated expression in a brain region (p<0.0001). There was also a significant overlap (>3 times greater than expected by chance) between isoforms from the same gene that had increased modification rates and longer polyA lengths within all brain regions (p<0.01). Therefore, when the same gene encodes multiple isoforms, those with distal polyadenylation sites (longer 3’ UTRs) will have increased m6A modification rates and longer polyA tails compared with other isoforms of the same gene. We also found that the number of m6A sites on an isoform was moderately correlated with polyA tail length (rho=0.1538, p<0.0001). The observed associations between polyA lengths, isoform expression, and m6A modification levels highlight the complexity of gene regulation and suggest that the interplay of polyadenylation and m6A modification patterns contribute to regulating gene expression in the brain.

## DISCUSSION

Our study is the first to apply direct RNA sequencing (DRS) on post-mortem human brain samples and we provide a transcriptome-wide map of >50k isoform-level m6A modification sites in three distinct brain regions. We identified widespread differences in isoform expression, modification and polyadenylation profiles between the brain regions. Notably, our findings were consistent across distinct analyses, with expression patterns of m6A machinery and isoform architecture reflecting the m6A modification levels in the different brain regions.

The highest and lowest proportion of m6A modified isoforms and highest and lowest modification rates at m6A sites were seen in CB and CN, respectively. Consistent with this finding, m6A writers displayed increased expression in CB and m6A erasers had increased expression in CN. We also found that isoform lengths and 3’ UTR lengths were longer in isoforms with DE in CB, and shorter in isoforms with DE in CN. Together, these results suggest that changes in the expression of the m6A machinery and tissue-specific isoform expression patterns are responsible for much of the overall differences in m6A levels between the brain regions.

While we found that most m6A modified isoforms were commonly modified in multiple brain regions, many were specifically modified in a single brain region and a majority of these were not due to region-specific upregulation of isoform expression. Although CB had the largest number of specifically modified isoforms, this region also had the largest proportion that were likely due to increases in expression. The PFC, in contrast, exhibited a distinctly high proportion of region-specifically modified isoforms that were not due to expression differences, underscoring the unique modification patterns in the PFC. The gene isoforms that displayed region-specific modification were associated with distinct cell types across brain regions. This suggests that region-specific modification may be due to cell-type-specific regulation of these isoforms. Isoforms specifically modified in CB were associated with cerebellar granule cells (CGCs) and it has been demonstrated that m6A is essential in this cell type during cerebellar development (59).

We also identified a set of hypermodified isoforms and many of these were consistently found in all brain regions. Hypermodified isoforms were enriched for excitatory neurons in all three brain regions and highly associated with multiple synapse and post-synapse GO terms, suggesting that regulation of the m6A modification profiles of these isoforms is involved in excitatory neuron function across many brain regions. Our study further emphasises the region-specific regulatory roles of m6A modification within distinct cellular contexts in the brain.

Due to the isoform-level resolution of m6A sites in our data, we were able to identify changes in modification rates between the same genomic site encoded in multiple isoforms. Generally, isoforms exhibited similar modification rates at these shared genomic sites and only ∼7% displayed differential modification (DM) rates. Consistent with recent studies, we found that DM between these isoforms was largely attributable to changes in the m6A site’s proximity to exon boundaries (33,40). We hypothesise that the small degree of DM in our study is a lower bound and increases in read depth and future improvements to m6A detection software will likely enable quantification of modification rates at more sites. Our results suggest that results from methods that provide peaks of m6A across gene bodies have likely masked many isoform-specific regulation events of m6A deposition.

We also identified thousands of m6A modified isoforms with differences in modification rates (DM) between brain regions, and most of these changes reflected the variable expression levels of m6A writers, readers and erasers. The isoforms with DM were significantly enriched for RNA-binding protein (RBP) sites and were also associated with multiple neuronal GO terms, as well as protein transport and microtubule polymerisation. Future research into whether these isoform- and tissue-specific m6A modifications are a result of distinct cell types or phenotypic states will be an important next step. The identification of m6A modifications in single-cells is an emerging field and is quickly becoming an area of great interest, however, these methods are still limited by the constraints of short-read sequencing (60,61).

Notably, synapse-related pathways were consistently associated with genes in various analyses in our study. The enrichment of synaptic terms in genes with differential modification and polyA lengths between brain regions underscores the significance of region-specific regulatory processes in synaptic functionality. Additionally, the association of synaptic GO terms in genes with differential isoform usage between brain regions implies that the expression of specific isoforms is important for modulating synapse activity in a region-dependent manner. Understanding synaptic regulation in various brain regions will be important in uncovering the mechanisms behind many neurological disorders linked to synaptic dysfunction (62).

A general critique of current m6A detection techniques, including immunoprecipitation-based methods, is that the results are not reproducible and several previous studies have lacked sufficient replicates (54). We aimed to address this by including at least three sample replicates per brain region, however, we note the moderate percentage (45%) of m6A sites detected in ≥3 samples per brain region. The number of reads and modification rates at these highly reproducible sites was significantly increased compared with the sites detected in ≥2 samples. Consequently, improving the read depth obtained per replicate will likely increase reproducibility at m6A sites. However, while recent benchmarking studies have shown that m6anet and xPore perform well, these programs may be limited in their ability to consistently detect m6A sites with low modification rates (55,56).

A current limitation of DRS is the relatively large sample input required and low number of reads generated, meaning the technique is not always feasible when only a small amount of sample or tissue is available for RNA isolation. The recent release of updated DRS kits from ONT aims to address the latter of these challenges and as sequencing throughput for DRS improves, there will be greater power to consistently detect m6A sites with low-medium coverage between replicates. Novel basecallers have also recently been introduced that aim to increase read accuracy and improve the single-molecule resolution at m6A sites (63).

Most gene-level m6A sites identified in our data were previously annotated in human tissues in the *m6A-Atlas* or *DirectRMDB* databases. We noted that m6A sites in lncRNAs represented a higher proportion of unannotated sites compared with annotated sites in our data and sites annotated only in *m6A-Atlas*, however not with the DRS-specific resource *DirectRMDB*. This suggests that DRS may be beneficial for investigating m6A within lncRNAs compared to previous methods, which will be particularly advantageous for profiling lncRNAs in the human brain, where they have integral roles in learning and memory (57,58). We also found that the PFC had the highest proportion of both unannotated m6A sites and genes found with only unannotated sites of the three brain regions. This result may be due to the additional replicate in PFC (n=4) compared to CB and CN (n=3). However, the total number of m6A sites identified was highest in CB, and CN had the second highest proportion of genes with only unannotated sites as well as the lowest number of m6A sites in total. While it is likely that there was increased power to detect more sites in the PFC, it is also possible that genes in this brain region have not been adequately profiled for m6A previously.

In summary, our findings have revealed new isoform-level insights into three distinct human brain regions. We have demonstrated the interplay of multiple RNA regulatory mechanisms such as isoform expression, m6A modification and polyadenylation. We suggest researchers move towards understanding the functional implications of m6A modifications in an isoform-specific and tissue-specific context and our study supports continued integration of long-read sequencing technologies into the field of RNA modifications.

## METHODS

### Sample preparation and quality control

Post-mortem brain tissue was obtained from six donors with no diagnosis or physiological evidence of neurological or neuropsychiatric disorders through the Victorian Brain Bank (VBB) under HREC approval #12457. The age, sex, post-mortem interval (PMI), brain tissue pH and brain weight for each individual are shown in Supplementary Table 1. Briefly, samples comprised both males and females (n=3 each), aged between 64-81 years, with PMIs between 24-59 hours. Frozen brain tissue was cut from three brain regions including prefrontal cortex (Brodmann’s area 46) (PFC), caudate nucleus (CN) and cerebellum (CB). Total RNA was extracted from bulk tissue across five randomised batches. Frozen brain tissue was homogenised on ice, using a manual tissue grinder (Potter-Elvehjem, PTFE), whilst immersed in 1 mL QIAzol Lysis Reagent (QIAGEN). The resulting lysate was then made up to 3 mL with QIAzol Lysis Reagent and mixed thoroughly before 1 mL lysate aliquots were processed using RNeasy Lipid Tissue Kit 74804 (QIAGEN) according to the manufacturer’s instructions. The increased volume of QIAzol Lysis Reagent was to ensure each RNA extraction column did not exceed the stated maximum binding capacity of approximately 100 mg. Three RNA elutions of 30 µL each were combined for a total of 90 µL for each sample. RNA quantity and quality were checked using a Qubit 4 Fluorometer (1 µL), TapeStation 4200 and Nanodrop 2000.

### Library preparation and direct RNA sequencing

Only samples with RINs >7 were selected for long-read direct RNA sequencing (DRS), as lower quality RNA was unlikely to yield informative results (64). There were 10 high-quality samples for DRS: 4 prefrontal cortex, 3 caudate nucleus and 3 cerebellum. Libraries were prepared on the same day where possible to reduce inter-run variability. PolyA+ RNA was isolated using NEXTFLEX® Poly(A) Beads (Perkin Elmer, NOVA-512980) with total RNA inputs ranging from 57-100 mg. Isolated polyA+ RNA (range: 350-500 ng) was used for library preparation with the direct RNA sequencing kit SQK-RNA002 (ONT). Spike-In RNA Variant (SIRVs) Isoform Mix E1 (Lexogen, 025.03) was added to the library at approximately 1% (5 ng) of the expected sample polyA RNA yield. Prepared libraries were sequenced on the ONT PromethION instrument using FLO-PRO002 flow cells and basecalled with Guppy (v6.0.17) to produce FASTQ files.

### Read alignment and quantification

Pass reads (qscore>7) in the FASTQ files were aligned to the human (GRCh38) and SIRV genome and transcriptome using minimap2 (v2.22). Genome alignments were performed using the splice-aware mode of minimap2 *-ax splice -uf -k14* and transcriptome alignments (GENCODE v31, SIRV) were performed using the long-read mode for ONT data *-ax map-ont*. FeatureCounts (v1.6.5) was used to quantify human and SIRV genome alignments with the parameters *-L – primary* to generate gene counts (65). NanoCount was used to quantify human and SIRV transcriptome alignments (v1.1.0) with default parameters to generate isoform counts (66). The BamSlam R script was used to obtain summary information regarding the transcriptome alignments outlined in Table 1 (66).

### Differential expression and isoform usage analysis

We used limma in R to test for differential gene and isoform expression between the three brain regions (67). Log2 fold changes and adjusted p-values (Benjamini-Hochberg) were calculated using the ‘voomWithQualityWeights’ function to account for any variation in sample quality and adjusted p-values <0.05 were required for significance (68). Differential isoform usage (DIU) analysis was performed in R using IsoformSwitchAnalyzeR (69). The isoform counts from NanoCount were input along with the annotation and transcriptome files. Statistical analysis was performed with DEXSeq to identify differential isoform usage between brain regions (70). The count matrix was filtered for genes with >1 isoform, genes with >20 counts and isoforms with >5 counts. We required a change in isoform proportions between brain regions of >0.2 to further increase stringency and an FDR-adjusted p-value of <0.05 for significance (71).

### Transcriptome-wide m6A modification sites

We used m6anet (v2.0.1) to identify N6-methyladenosine (m6A) sites in DRACH motifs (D: A, G or U, R: A or G, H: A, C or U) from our direct RNA reads in each sample. The program outputs an m6A modification probability and modification rate (proportion of modified reads) at every transcript isoform site with a coverage of >20 reads (18). We required a modification probability of >0.90 in >1 sample for the site to be classed as m6A modified (n=57,144 unique m6A sites, n=228,314 total m6A sites across all samples).

### Identification of common and brain-region-specific m6A isoform modifications

Commonly modified genes/isoforms had at least one m6A modification site identified in all three brain regions. To identify the features with brain-region-specific modification, we subset genes and isoforms to those only found with m6A modifications in a single brain region. To remove the potential effect of brain-region-specific expression causing genes or isoforms to have specific modification, we removed those with specific expression in only a single brain region as found in the differential expression analysis. The common and brain-region-specific m6A modified genes were used in a gene ontology (GO) and cell-type-specific enrichment analysis. The distributions of common or brain-region-specific m6A modifications along isoforms were compared using a Kolgomorov-Smirnov test (Figure 5C).

### Comparison with annotated m6A sites in *m6A-Atlas* and *DirectRMDB*

We downloaded data from *m6A-Atlas* (v2.0) and *DirectRMDB* to identify genomic m6A sites in our data that were previously annotated (27,28). The data was subset for only human cell lines or tissues for comparison.

### Hypermodified and unmodified isoforms

We calculated a normalised number of m6A modifications per isoform by summing the modification rates at every m6A site along the isoform and using this along with isoform length and exon density in a linear regression as predictor variables. We extracted the residuals from this model and used these as normalised m6A values. We defined the hypermodified isoforms as the top 500 per brain region ranked by normalised m6A values. The unmodified isoforms were defined as those with no m6A modifications that also had adequate coverage for m6A detection (>1 DRACH motif detected in m6anet) and a modification probability of <0.5 at all detected sites within the isoform.

### Identification of differential modification between isoforms and brain regions

We used a two-proportion z-test to identify differential modification (DM) rates between isoforms encoding the same genomic m6A site within a brain region. We required an FDR-adjusted p-value <0.05 and a modification rate difference between isoforms of >0.15 for significance.

To test for differential modification at the same site in an isoform between different brain regions, we integrated the results from m6anet and xPore (41). xPore (v2.1) identifies sites with differential modification rates between conditions but does not identify the type of modification present when it is run without an unmodified control sample. We subset the xPore sites for DRACH motifs, a modification rate difference of >0.3 and an FDR-adjusted p-value<0.05 as recommended (41). These sites were then overlapped with sites found in m6anet (modification probability score >0.7) in >1 sample to create the final list of differentially modified sites.

### PolyA tail length quantification and analysis

The polyA tail lengths for each read were estimated using the ‘polya’ module from nanopolish (v0.13.2) (53). We kept reads assigned a ‘pass’ QC tag from nanopolish and required >5 reads per isoform per sample. Consistent with previous studies, we found that mitochondrial isoforms had shorter polyA tail lengths than non-mitochondrial isoforms, so these were excluded from downstream analysis (53). We calculated a median polyA tail length for each isoform in each sample to avoid highly expressed isoforms skewing the comparisons. We performed a Mann-Whitney-Wilcoxon (MWU) test to compare the overall median polyA lengths between the brain regions and between isoforms of the same gene. We used a MWU test for this comparison and required >50 reads and a polyA length difference of >20 nt per isoform.

We noted that some isoforms had large variations in their estimated polyA lengths, termed ‘dynamic’ polyA tails. We used the interquartile range (IQR) of isoforms with >50 reads per brain region to rank the top 250 isoforms according to variations in polyA lengths per brain region. IQR was used to prevent a small number of outliers from influencing the ranking.

### Data analysis and visualisation

We used R for all statistical analysis and plotting unless otherwise stated. The MWU test was used for statistical comparisons with the wilcox.test() function from the ‘stats’ package and p-values were subsequently FDR-adjusted when multiple comparisons were performed with the p.adjust() function also from the ‘stats’ package. Linear regressions were performed using the lm() function and their respective summaries were extracted using the summary() function. The metagene plots were produced using metaPlotR (72). Hypergeometric tests were performed with phyper to obtain p-values for the number of overlapping isoforms between different analyses.

We used the R package clusterProfiler (v4.6.2) to perform the various gene ontology (GO) analyses in this study (73). The enrichGO() function was applied to gene sets using relevant background genes (i.e. expressed genes, m6A modified genes). All three GO domains were included: biological process, molecular function and cellular compartment. P-values were adjusted for the false-discovery rate (adjusted p-value<0.05) and redundant GO terms were removed using the ‘simplify’ function with a cutoff value of 0.7. We removed GO terms with <10 genes assigned to the pathway and also calculated an enrichment value for each GO term as per ShinyGO (v0.77) defined as the percentage of genes belonging to a pathway divided by the corresponding percentage of background genes belonging to the pathway (74).

To perform the cell-type-specific enrichment analysis we used WebCSEA (32) and required a combined p-value <0.001 for significance and subset the results for the ‘adult’ development stage and ‘nervous system’ organ system as recommended.

Data for all annotated human RNA binding protein sites was downloaded from POSTAR3 and CLIPdb (43,44). The sites were filtered for those annotated at least two times. We used bedtools to intersect the RNA binding protein sites with genomic coordinates of all DRACH sites tested with m6anet, all m6A sites identified by m6anet, and m6A sites with differential modification rates between brain regions identified by xPore, with the following command: bedtools window -a xPore_genomic_positions.bed -b human_RBP_sites.bed -w 5 -u > result.bed. We used the two-proportion z-test to determine if there were significant differences in proportions of sites overlapping RBP sites (Figure 7C,D).

## ACKNOWLEDGEMENTS

The authors would like to thank the Leichtung family for funding the Brain and Behavior Research Foundation NARSAD grant, as well as members of the Clark lab for useful feedback on the manuscript. The authors would also like to thank Geoff Pavey and Fairlie Hinton (VBB) for their assistance with frozen tissue selection and preparation, Sarah MacRaild and Dr Quentin Gouil (WEHI) for their assistance with Nanopore PromethION sequencing. This research was undertaken using the LIEF HPC-GPGPU facility hosted as part of Spartan at the University of Melbourne. This facility was established with the assistance of LIEF Grant LE170100200.

The authors would like to acknowledge that brain tissues were received from the Victorian Brain Bank, supported by The Florey, The Alfred and the Victorian Institute of Forensic Medicine and funded in part by Parkinson’s Victoria, MND Victoria and FightMND. We further acknowledge the donors and their families for their selfless donations to research.

## AUTHOR CONTRIBUTIONS

J.G designed and performed the bioinformatic analysis, interpreted results and wrote the manuscript with assistance from all authors. R.D.P designed and performed experiments, wrote and reviewed the final manuscript. M.B.C designed experiments, supervised research, and wrote and reviewed the final manuscript. C.M classified and selected brain samples in conjunction with the VBB. S.U.M and T.W.B provided assistance with data analysis and interpretation, and reviewed the final manuscript.

## CODE AVAILABILITY

https://github.com/josiegleeson/BamSlam https://github.com/josiegleeson/annotate-m6anet-output https://github.com/josiegleeson/compare-modification-rates-at-shared-sites

