## Supplementary Figures for "Isoform-level profiling of m6A epitranscriptomic signatures in human brain"

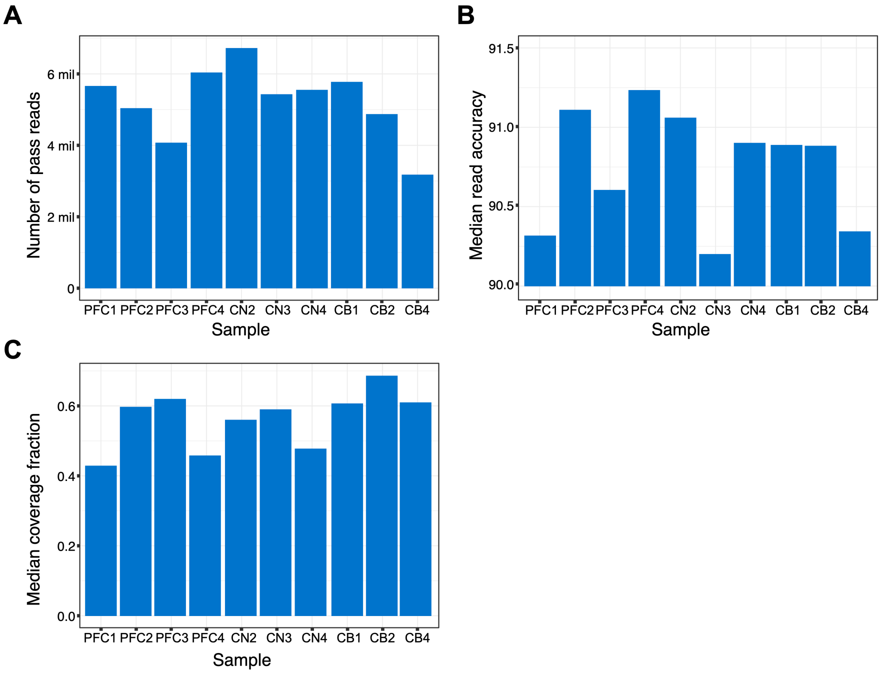


**Supplementary Figure 1. Metrics for DRS of 10 samples (from BamSlam).**
**(A)** Total number of pass reads per sample. **(B)** Median read accuracy per sample calculated from CIGAR strings in BAM files as: (M+I+D-NM)/(M+I+D). **(C)** Median coverage fraction of reads per sample calculated as: alignment length / mapped isoform length.

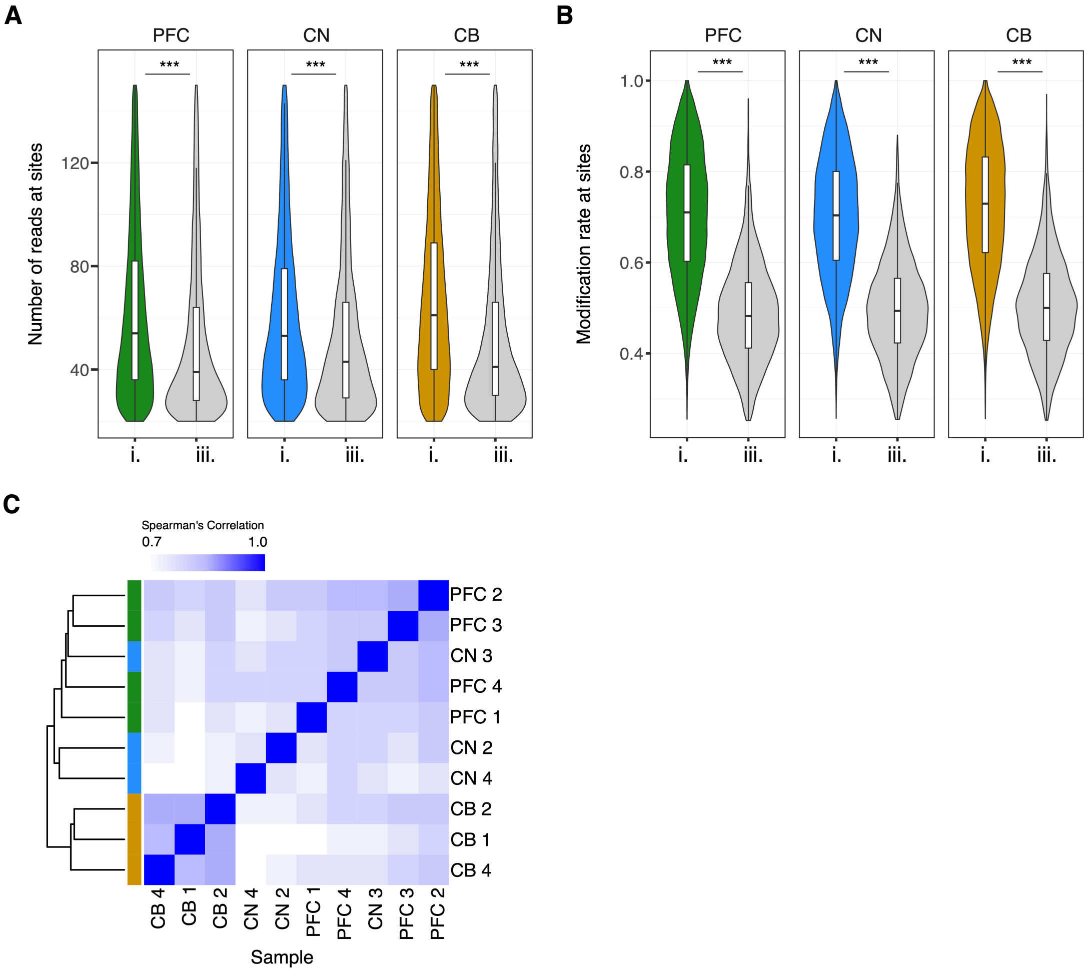


**Supplementary Figure 2. Read coverages and modification rates at m6A sites.**
**(A)** The number of reads at m6A sites and **(B)** modification rates at m6A sites, detected in i. ≤2 samples per brain region or iii. ≥3 samples per brain region (i. and iii. are consistent with Figure 4D). **(C)** Heatmap of correlations between modification rates of m6A sites per sample.

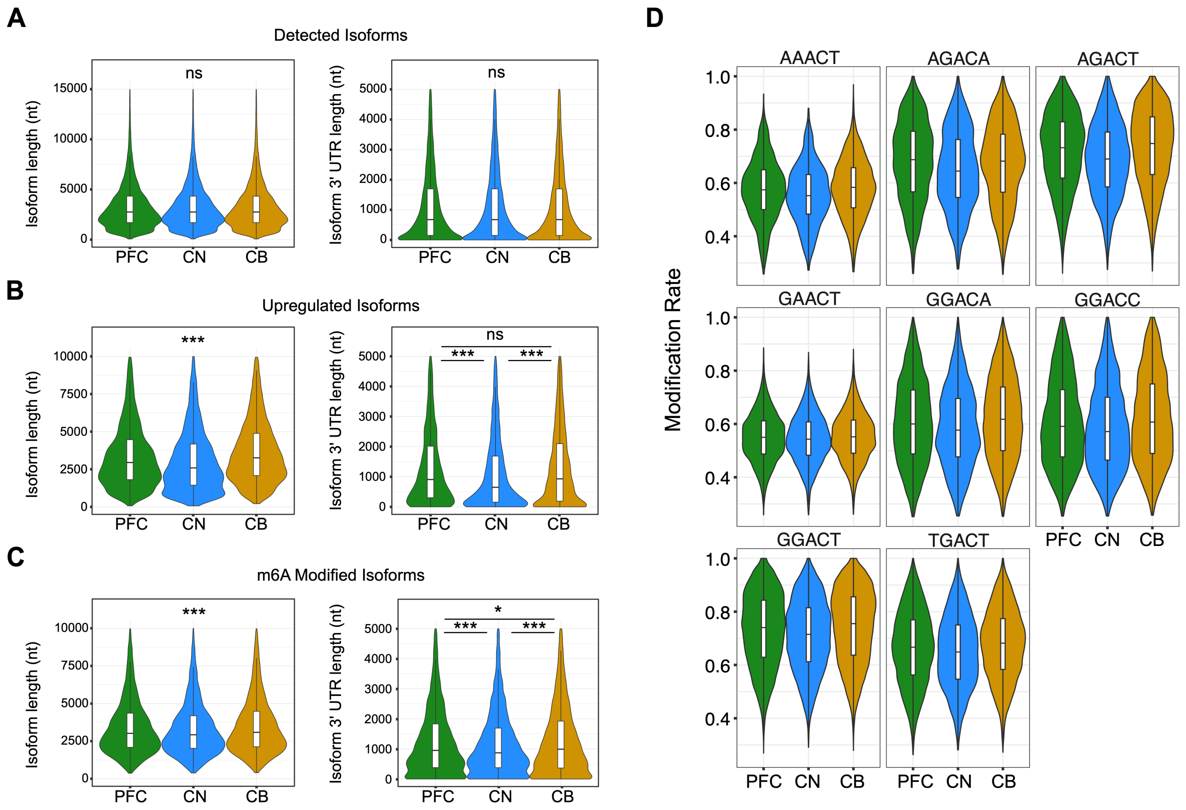


**Supplementary Figure 3. Isoform lengths, 3’UTR lengths and modification rates in each brain region.**

The isoform lengths (left plots in panels A-C) and 3’ UTR lengths (right plots in panels A-C) were compared between brain regions for different isoform sets. **(A)** Detected isoforms (counts >5) showed no significant differences between brain regions. **(B)** Isoforms that were upregulated (DEIs) in their respective brain regions showed significant differences in length between all brain regions, and significantly longer 3’UTRs in PFC and CB compared to CN. **(C)** Isoforms that were m6A modified in each of the respective brain regions showed significant differences in both isoform length and 3’UTR length between all brain regions. P-values from Mann-Whitney-Wilcoxon tests are indicated by ‘ns’ for >0.05, ‘*’ for <0.05 and ‘***’ for <0.001, where only one value is shown, all comparisons had this consistent value. **(D)** Modification rates at m6A sites plotted for each DRACH motif per brain region.

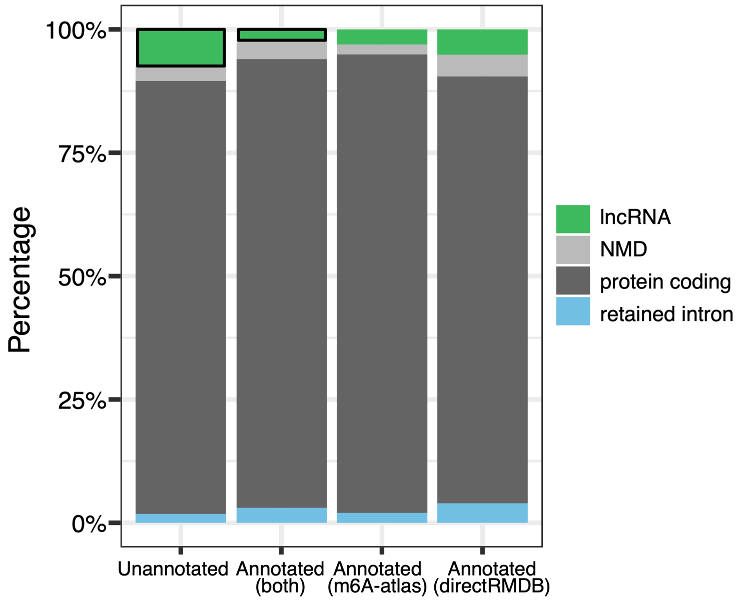


**Supplementary Figure 4. LncRNAs in unannotated vs annotated m6A sites.**
A higher percentage of previously unannotated m6A sites are within lncRNAs compared with those annotated in both *DirectRMDB* and/or *m6A-Atlas* (boxes with black outlines indicate lncRNA categories in each group). Unannotated lncRNA = 7.4%, annotated in both databases lncRNA = 2.2%, p-value<0.0001 (two-proportion z-test).

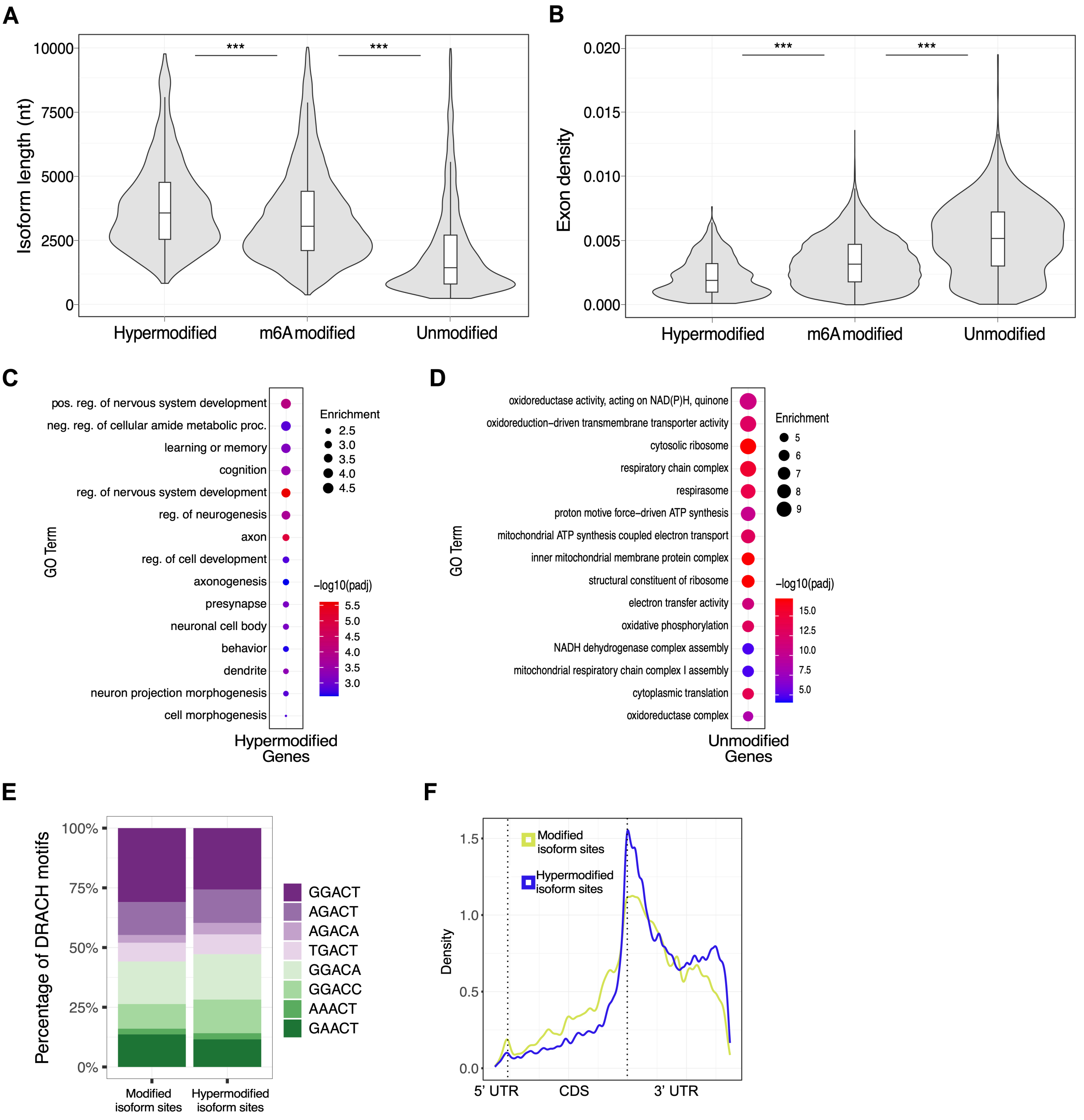


**Supplementary Figure 5. Comparison of hypermodified, modified and unmodified isoforms.**

**(A,B)** Hypermodified, modified and unmodified isoforms were compared for **(A)** isoform length (nt) and **(B)** exon density per isoform (exon density = number of exons / isoform length). Significance of MWU p-values is indicated by '***' for <0.001. **(C)** GO analysis of genes from 911 hypermodified isoforms. (**D)** GO analysis of genes from 3,907 unmodified isoforms. **(E)** Percentages of each DRACH motif in all m6A sites and m6A sites within hypermodified isoforms. **(F)** Metagene plot showing the distribution of all m6A sites and m6A sites in hypermodified isoforms (Kolmogorov–Smirnov p<0.0001).

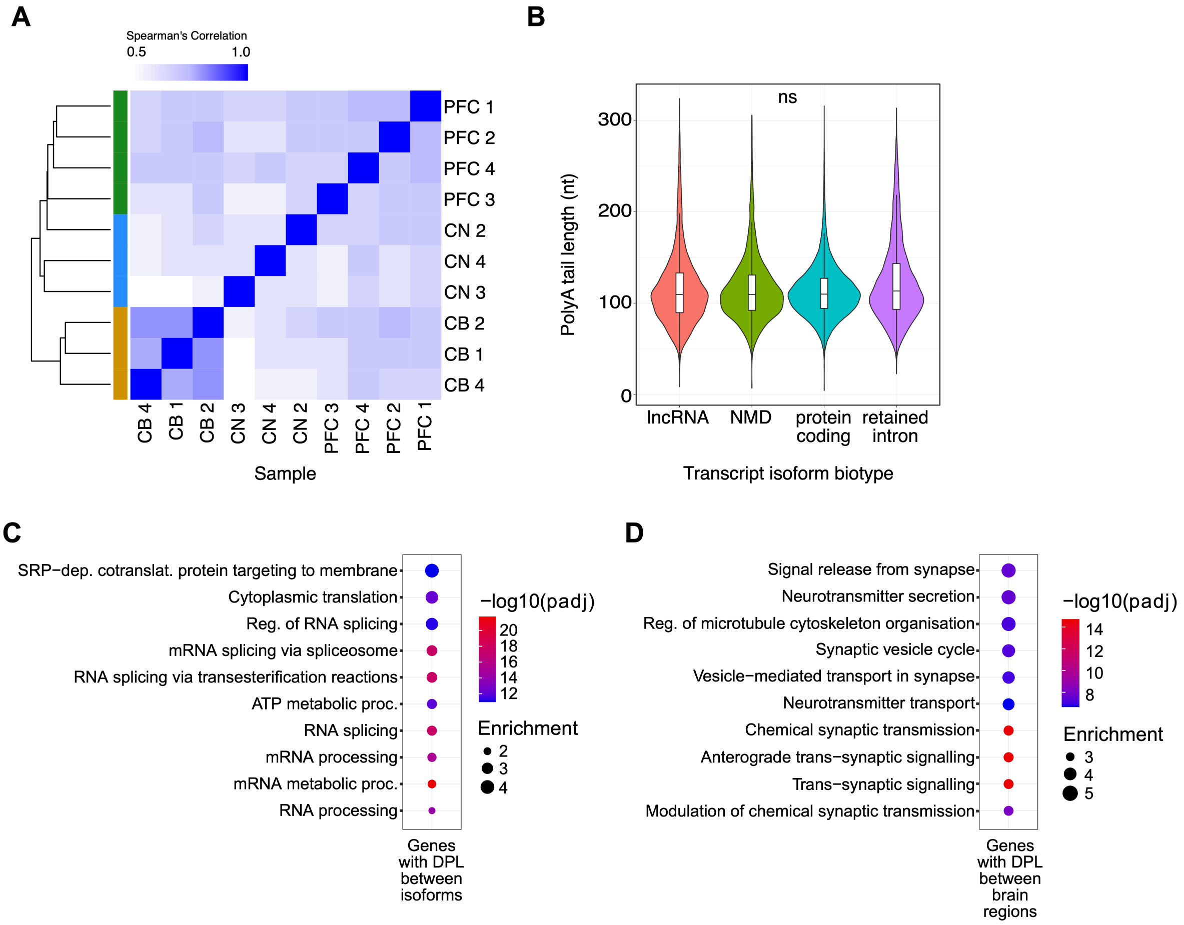


**Supplementary Figure 6. PolyA length comparisons and GO analysis.**
**(A)** Heatmap showing that samples from the same brain region largely clustered together based on the median polyA length per isoform. **(B)** Violin plots of polyA lengths between different transcript isoform biotypes. No significant differences in polyA lengths were observed. **(C,D)** GO analysis of genes with **(C)** differential polyA lengths (DPL) between isoforms in a single brain region and **(D)** DPL between the same isoform in different brain regions.
